# A modelling challenge to improve preparedness for highly pathogenic avian influenza epidemics: objectives, design and synthetic data generation

**DOI:** 10.64898/2026.10.06.756671

**Authors:** Brandon Hayes, Gaël Beaunée, Sébastien Lambert, Clara Delecroix, Timothée Vergne

**Author notes:** **Author for correspondence:** Brandon Hayes,.

## Abstract

Highly pathogenic avian influenza (HPAI) is among the most significant disease threats to poultry worldwide. Modelling challenges, where independent teams address a common epidemiological scenario, can support the development of response capabilities and strengthen communication between modellers and risk managers. Despite the established benefits of these exercises, they remain uncommon in animal health, and none have previously addressed avian influenza. Here Jolly Island, an artificial landscape of 9160 poultry farms with five production types, was developed and seeded with stochastic HPAI epidemics that ran their course over a three-to-four month period. From January through April 2026, 45 registered teams across five continents investigated a selected synthetic epidemic, producing probabilistic forecasts, inferring transmission drivers, and evaluating counterfactual control policies. Here the objectives, design choices, and data generation of the HPAI Modelling Challenge are described in full, covering transmission processes, surveillance system operations, and control strategy implementation, along with the selected epidemic trajectory, the narrative situation reports supplied to the teams at each phase, and the complete phase-specific datasets released to them. As the opening paper of this special issue, here is provided the foundation upon which all subsequent contributions, where teams document their approaches and analyses, are built.

## 1. Introduction

Mathematical and statistical models are increasingly used to support the decision process in infectious disease outbreak response (World Health Organization et al., 2024). In their employ, these computational tools help overcome the difficulty of projecting likely outcomes of alternative response policies in a rapidly changing environment. Modelling has repeatedly proven its utility, including during the responses to pandemic H1N1 (Lee et al., 2013), Ebola (Meltzer et al., 2016), and COVID-19 (Howerton et al., 2023) in human health, and to foot-and-mouth disease (Keeling et al., 2001), H7N7 avian influenza (Stegeman et al., 2004), and African swine fever (Pepin et al., 2020) in animal health. The operational value of these models, however, depends not only on predictive accuracy, but also on the ability to deploy them in a timely manner, integrate disparate data sources, characterise uncertainty, evaluate interventions, and effectively communicate results to decision makers (Rivers et al., 2019).

Modelling challenges provide a means to develop, test and compare the capabilities of these mathematical and statistical tools in realistic response conditions. These friendly challenges consist of inviting independent teams to address a common predefined problem, and in doing so contribute to both development of modelling capacity and improvement in communication between modellers and risk managers (Rivers et al., 2019; Sherratt et al., 2025)

Whereas this approach is well-established in human health, as illustrated by the hubverse, a recently-developed open-source infrastructure to facilitate collaborative modelling (Kerr et al., 2026), in animal health it remains nascent with only a single exercise having been conducted to date: The African swine fever modelling challenge (Ezanno et al., 2023). Over a six-month period from August 2020 to January 2021, independent teams were tasked with characterising the drivers of an African swine fever (ASF) epidemic, forecasting spatiotemporal infection patterns, and evaluating candidate control measures (Picault et al., 2022). Overall, the event was found to have improved the readiness of the modelling community for future ASF outbreaks, and supported the use of ensemble rather than individual models in addressing policy-relevant questions for ASF response (Ezanno et al., 2022). Extending this approach to other major infectious diseases in the animal health field would further help reinforce modelling capabilities and international preparedness.

Highly pathogenic avian influenza (HPAI) is an excellent candidate for such an extension, as it represents one of the most significant infectious disease threats to poultry globally (FAO and WOAH, 2025). Successive waves of HPAI H5 clade 2.3.4.4b in poultry have caused unprecedented mortality across Europe (European Food Safety Authority (EFSA) et al., 2026), (Kondo et al., 2025), and the Americas (Congressional Research Service, 2025), with immense consequences for trade (Padilla et al., 2025), food security (European Food Safety Authority (EFSA) et al., 2026), and animal welfare (EFSA Panel on Animal Health and Welfare (AHAW) et al., 2019). Both its persistence in wild bird populations (Couty et al., 2026) along with a growing capacity to adapt to mammals (including humans) (Capelastegui and Goldhill, 2025) has elevated HPAI to a perennial threat in much of the world. Sustained global preparedness is essential to mitigate its impacts (Kojima et al., 2024). However, the modelling effort itself has been directed almost exclusively to that of retrospective analyses of past epidemics, rather than operational, real-time support during outbreaks (Lambert et al., 2023).

The HPAI modelling challenge (hereafter referred to as the Challenge) was therefore developed to address this gap, with the objectives of strengthening modelling readiness for HPAI and fostering international and interdisciplinary collaboration. Over the course of a simulated four-month epidemic, participating teams were tasked with evaluating and synthesizing surveillance data, generating probabilistic forecasts, producing scenario projections, and providing evidence-based policy recommendations. In this article, we describe the objectives and design of the Challenge, along with the stochastic agent-based model used to generate the synthetic data provided to participating teams.

## 2. METHODS

### 2.1 Challenge setup and characteristics

Challenge preparation consisted of defining objectives, preparing the data to be sent to participating teams, designing the questions asked, and recruiting teams.

Two primary objectives were defined for the Challenge: to strengthen operational modelling readiness for HPAI, and to foster international and interdisciplinary collaboration both among modelling teams and between modellers and risk managers. To better meet these objectives, inference results, forecasts, and policy recommendations from participating teams needed to be able to be compared to an underlying ground truth, and intervention scenario alternatives needed an observable counterpart. As real outbreak data would neither provide such a foundation nor be unconstrained by confidentiality issues, the Challenge was built on fully synthetic data from a generative model described herein. The focus was maintained on the poultry sector, where the burden of HPAI is high and where decision-support modelling would be of the greatest relevance.

Questions were designed to span four analytical categories of increasing difficulty: descriptive epidemiology, short-term forecasting, transmission inference, and counterfactual policy evaluation. Descriptive and forecasting tasks were made so as to be accessible to all teams regardless of approach, while the inferential and counterfactual tasks were expected to require mechanistic methods. Such construction allowed teams of varied methodological backgrounds to participate, while still assessing a wide range of capabilities relevant to outbreak response. Descriptive epidemiology questions were mandatory, while all other questions were facultative, leaving each team with the choice to answer depending on their methods, time, and expertise.

To guide scenario design and to ensure policy relevance of Challenge objectives, consultations were held with risk man-agers and veterinary authorities from the onset. Controlling HPAI in poultry requires rapid detection and depopulation of both infected and high-risk farms, along with movement restrictions and heightened surveillance activity in established regulatory zones. The constraints that would have the greatest effect on the success of this response were therefore identified during these consultations and prioritised for incorporation into the modelling framework: culling capacity limitations, balancing reactive and preventive culling prioritisations, zone-based application of surveillance and control, and accounting for unobserved transmission pathways (e.g. wildlife, shared equipment and personnel).

To recruit modelling teams, the Challenge was advertised through a variety of avenues: on the project website, by email, and on professional social networks. Participation to the Challenge was open to any team irrespective of disciplinary background or prior modelling experience, and was a deliberate choice to encourage a heterogeneous set of analytical approaches.

To ease data familiarisation and facilitate the development of analytical pipelines, teams were provided with population data and a dummy case series one month prior to Challenge launch. A virtual kick-off meeting was organised on 29 January 2026 to introduce the Challenge scenario, datasets, and questions.

Finally, the Challenge itself (during which participating teams received data and submitted analyses) ran from 29 January to 30 April 2026, and was structured into three phases approximately one month each, coinciding with three epidemic periods of onset, peak, and resolution (Table 1). Two timescales are therefore used throughout: a phase is a working window for the teams; a period is a stage of the synthetic epidemic. Each phase was aligned to its corresponding period by design, so that Phase 1 tracked the epidemic through the end of Period 1, Phase 2 through the end of Period 2, and the same for Phase 3 with Period 3. However, phases and periods did not occur simultaneously, as the epidemic dates precede or fall early within the working window to which they correspond. This means that the data packages which were released at the start of each phase contained its period in full, and that teams were analysing an epidemic that had ran through the end of the period on which they were working.

**Table 1.** Alignment of player phases and epidemic periods. Phases are the working windows during which teams receive data packages and submit analyses. Periods are the stages of the synthetic epidemic that coincide with a phase. Day 0 is the day of first detection in the selected trajectory and when the epidemic became observable, 21 December 2025 (simulation day 8).

| Phase (teams) | Player working window | Epidemic period | Days (and in-simulation dates) from first detection |
| --- | --- | --- | --- |
| Phase 1 | 29 Jan 2026 – 28 Feb 2026 | Period 1 (onset) | –7 to 23 (21 Dec 2025 – 13 Jan 2026) |
| Phase 2 | 2 Mar 2026 – 31 Mar 2026 | Period 2 (peak) | 24 to 54 (14 Jan 2026 – 13 Feb 2026) |
| Phase 3 | 1 Apr 2026 – 30 Apr 2026 | Period 3 (resolution) | 55 to 111 (14 Feb 2026 – 11 Apr 2026) |

At the start of each phase, teams received a data package and a narrative situation report, containing both epidemic context details along with a question series requiring their response, and had four weeks to submit their analyses before the next data package was released. Each data package was right-censored at a fixed in-simulation date that marked the end of the corresponding period. Thus, a team working during Phase 1 (29 January to 28 February) analysed the epidemic as it was observed only up through 13 January, reproducing the lag between data accrual and analysis that characterises real outbreak response. Communication during the challenge between teams and with the organisers was maintained through a dedicated Slack workspace (Slack Technologies LLC, 2026).

### 2.2 Construction of the synthetic poultry system

#### 2.2.1 Jolly Island

A fictional island (“Jolly Island”) was created by merging all communes within the Pays de la Loire and Nouvelle-Aquitaine regions of France, then removing those from the Mayenne, Sarthe, and Pyrénées-Atlantiques departments (GADM, 2025). This landmass underwent an affine transformation in which it was reflected, rotated, and contracted around its centroid then translated to the mid-Atlantic just north of the Azores (Fig. 1), aligned to plausible waterfowl migratory flyways, yielding an island of approximately 35,000 square kilometres. CORINE landcover data (European Union, 2018) of the original landmass and a location-appropriate coordinate reference system (UTM 25N) were further incorporated. The final synthetic island contained 14 counties (renamed from departments), and 556 districts (renamed from cantons), having original names substituted with those of locations from New Jersey and Pennsylvania (out of first-author affinity and nostalgia).

**Figure 1.**
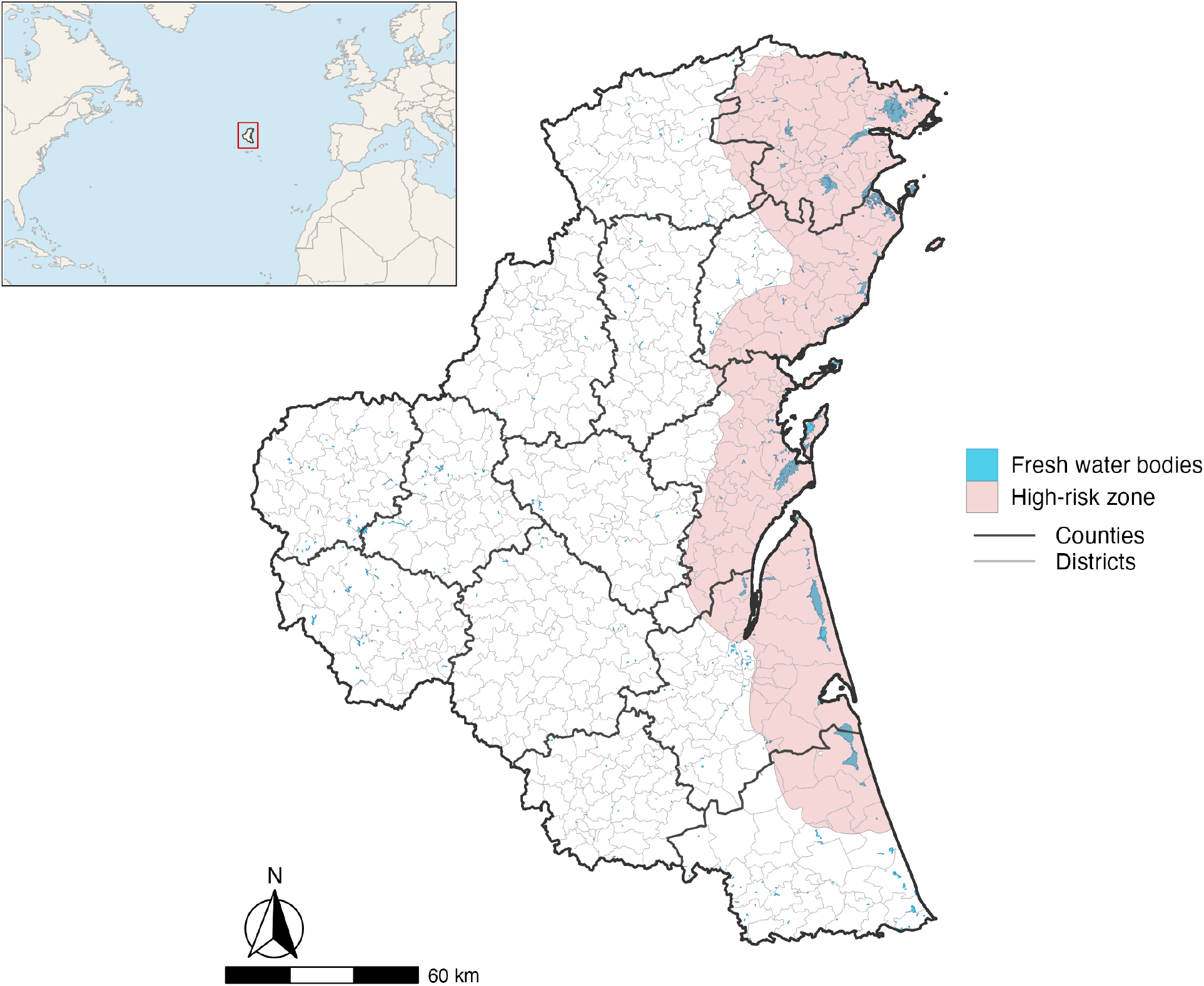
The synthetic Jolly Island (∼35,000 km^2^), situated in the mid-Atlantic just north of the Azores (inset), with freshwater body coverage and associated wild-bird high-risk zone. The 14 counties are shown in heavier outline, with the 556 districts distinguished by lighter outlines.

The poultry population comprised 9,160 farms, distributed among two species and five production types: chicken broiler stage 1 (*n* = 1, 248), chicken broiler stage 2 (*n* = 2, 746), chicken layer (*n* = 441), duck conventional (indoor, *n* = 3691), and duck organic (outdoor, *n* = 1, 034). Farm types were intentionally kept simple as mono-species operations. The two-stage broiler system was incorporated to simulate interconnected production cycling, where a larger stage 1 operation distributes chicks to multiple smaller stage 2 farms for finishing. Each farm contained attributes for species, production type, coordinates (see below), flock capacity, and a risk score computed from freshwater body proximity, which was used as a proxy for wild bird exposure. This risk scores were used to define the high-risk zone (Fig. 1) for farms close to water bodies, where active surveillance was performed.

Farms were distributed throughout the island via a structured stochastic process. The island contained two densely populated poultry areas (DPPA), one each in the north and south, where each one consisted of a core of connected districts plus a 12 km transition ring. Together, they contained 4,126 farms, with a density of 0.66 farms km-2 in the northern and 0.52 in the southern area, decreasing to 0.39 and 0.33 in the respective transition areas. The rest of the island had an average density of 0.21 km-2 (illustrated in Supplementary Material, Section 1, Figure S1). In the northern DPPA, duck farms were over-represented relative to chicken farms, and in the southern DPPA chicken farms were over-represented relative to duck farms. To ensure that control measures remained effective under realistic spatial constraints, local farm density distributions (i.e. the number of neighbouring farms within 1 km and 3 km radii) were calibrated to those observed in the French poultry production systems, having been the inspiration for the applied interventions, with the landmass resized accordingly. Conversely, farm production types and proportions, movement patterns, and spatial distribution were designed to encompass the types of heterogeneity seen in real production systems without replicating that of any specific country or region. Flock capacities were similarly assigned as a reflection of realistic commercial sizes, with each farm’s population drawn from a production-type-specific capacity distribution representative of those of a Western European context.

#### 2.2.2 Farm activity and movement

Farm production dynamics were incorporated through individual farm cyclic activity schedules, defined by a farm’s production type. Farms alternated between active and inactive periods, with activity defined as having birds present and permitting a farm to be susceptible to infection. Inactive periods corresponded to downtime periods, where the farm was cleaned and disinfected and kept empty for several days before restocking. Active production periods lasted approximately three weeks for stage 1 broiler farms, four weeks for stage 2, seven to eight weeks for conventional (indoor) duck farms, ten to twelve weeks for organic (outdoor) duck farms, and around thirteen months for layer farms, each followed by a minimum downtime of three weeks prior to restocking.

Layer, stage 1 broiler, and conventional and organic duck farms were assumed to receive all shipments that initiate a production cycle from HPAI-free off-island sources. Layer, stage 2 broiler, and conventional and organic duck farms were all assumed to send their flocks en-masse to abattoirs at the end of their production cycle (not explicitly simulated). Broiler farms relied on a coupled supply chain, in which stage 2 broiler farms became active on a day only if a shipment was received from a stage 1 broiler source on that day. Movement data between farms was used to generate farm activity calendars, with baseline shipment and activity schedules precomputed over a multi-year period encompassing the planned epidemic dates.

### 2.3 Model used to produce synthetic epidemiological data

To produce the synthetic epidemic, a stochastic farm-level model was developed with the expectation that it would be more complex than any model developed by participating teams. The model ran in discrete time using daily time steps.

Stochastic processes were evaluated at a subdaily resolution (Δt = 0.1 days), providing a time-discretised approximation to continuous-time processes as consistent with classical *τ* -leap methods (Keeling and Rohani, 2008). The simulation was implemented in R (R Core Team, 2024) with C++ integration of high-performance components via the Rcpp package (Eddelbuettel and Francois, 2011).

#### 2.3.1 Within-farm dynamics

A deterministic SEIRD (Susceptible-Exposed-Infectious-Recovered-Dead) compartmental model of within-farm dynamics (at the level of individual birds within a farm) was used to generate production-type-specific infection trajectories (Fig. 2). Production-type-specific parameters governed within-farm transmission rates and mortality-based suspicion thresholds, while species-specific parameters governed within-farm latent and infectious period durations and disease-induced mortality probability. The infectious period was modelled as Erlang distributed, a gamma distribution with an integer shape parameter, equivalent to a sequence of n exponential sub-stages, rather than exponentially. The rationale for this choice is detailed in Section 4. Smaller broiler and organic farms were assumed to follow density-dependent transmission while larger conventional duck and layer farms followed frequency-dependent transmission, reflecting the assumption that the number of effective contacts per bird increased with flock size in smaller farms but saturated in very large flocks.

**Figure 2.**
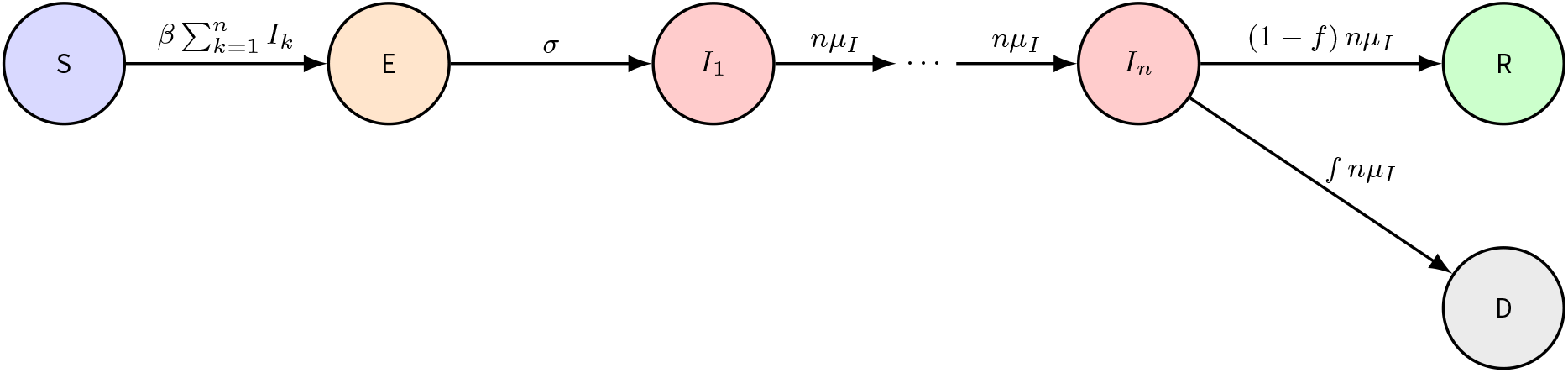
Within-farm SEIRD compartmental structure. The density-dependent formulation is shown; for production types following frequency-dependent transmission, the force of infection is given by 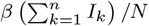 in place of 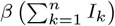 . Transition rates are defined as *σ* = 1*/*latent period, *µ*_*I*_ = 1*/*infectious period, and *f* = disease-induced mortality probability. The infectious compartment is represented as a chain of *n* sub-stages (*I*_1_, …, *I*_*n*_), resulting in an Erlang-distributed infectious period (the integer-shape special case of the gamma distribution), such that total infectiousness is given by 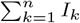 .

The within-farm model was solved once per production type prior to initiating the between-farm stochastic simulations (see Supplementary Information, Section 2 for full parameterisation). Model outputs were production-type-specific and consisted of the time from infection to infectiousness, first-death, and suspicion, along with the prevalence trajectory for the median-sized farm of that production type. Event times are provided in Supplementary Information, Section 2, Table S5.

Within-farm infection dynamics were mapped to between-farm transmission via a function of the time since a farm became infectious. For each production type, the outwardly radiating infectious pressure was weighted as the prevalence of infectious animals over time, resulting in a between-farm infectious intensity that followed the within-farm infectious profile.

#### 2.3.2 Between-farm transmission

Between-farm transmission occurred via three routes: community (local), network (shipment-mediated), and wildlife exposure (external). In community transmission, each susceptible recipient farm *j* accumulated a force of infection (FOI, *λ*) from all infectious farms *i* within the transmission distance limit, as given by:

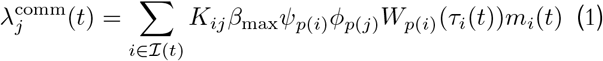

where (*t*) is the set of infectious farms on day *t, K*_*ij*_ is the spatial transmission kernel for infectious-susceptible farm pair *ij* (Eq. 2), *β*_max_ is the baseline maximum transmission rate for the community route, and *ψ*_*p*(*i*)_ and *ϕ*_*p*(*j*)_ are the production-typespecific infectivity and susceptibility modifiers, respectively (see Supplementary Information, Section 2 for parameterisation). Within-farm dynamics are linked to between-farm transmission intensity through the term *W*_*p*(*i*)_(*τ*_*i*_(*t*)), the proportion of birds that are infectious within a farm of production type *p*(*i*) and evaluated at *τ*_*i*_(*t*), the number of days since farm *i* became infectious. Lastly, *m*_*i*_(*t*) is an infectiousness multiplier reflecting on-farm biosecurity measures that are implemented once a farm is suspicious of being infected. It equals 1 for farms of unknown status and takes a lower value once a farm is suspected or detected. This reduction becomes stronger in successive epidemic periods as it is assumed both awareness and biosecurity intensity increase as the epidemic progresses (Supplementary Information, Table S9).

The spatial kernel took the three-parameter logistic expression form from Boender et al. (2007):

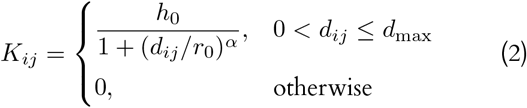

where *d*_*ij*_ is the distance (km) between farms *i* and *j, h*_0_ is the baseline kernel height, *r*_0_ is the scale parameter that gives the distance at which the kernel decreases to half of *h*_0_, and *α* is the shape parameter that governs the rate of decay. The kernel decreases with distance and was truncated to zero beyond *d*_max_ = 30 km.

Shipment-mediated transmission was implemented as a separate deterministic process through the broiler supply chain, from infected stage 1 farms to stage 2 destination farms. Given that there was an approximate average of 2200 birds per shipment (range 491–5175), and shipment bins were constructed through random draws from across the flock, infection in all shipments from a source was assumed certain if the source had a non-zero within-farm prevalence at the time of shipment. Planned shipments involving detected-as-infected source farms, recipient farms that had been banned from restocking, or zone-restricted farms were blocked.

External transmission, presumed to arise from exposure to infected wild birds, was computed as a stochastic infection hazard using a period-specific transmission rate:

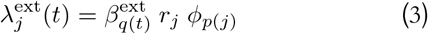

Where 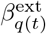 is the period-specific external transmission rate in epidemic period *q*(*t*) *∈*1, 2, 3, *r*_*j*_ is a farm-level relative risk score derived from proximity to freshwater bodies, and *ϕ*_*p*(*j*)_ is the production-type-specific susceptibility. The external transmission rate decreased across successive epidemic periods, until there was no transmission in the final period as migratory wild birds were assumed to have left the island (Supplementary Information, Section 2.3, Table S8).

The total FOI acting on farm *j* was therefore defined as the sum of the FOI of each of the two stochastic transmission routes:

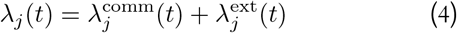

At each sub-step Δ*t*, the FOI was converted into an infection probability using the standard rate-to-probability conversion:

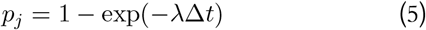

and infection was determined by a Bernoulli trial for each susceptible farm.

#### 2.3.3 Surveillance and detection

Infected farms could be detected through several methods: passive surveillance, enhanced passive surveillance, pre-shipment testing, contact tracing or as they underwent preventive culling. Passive surveillance occurred through a production-type-specific mortality-driven suspicion trigger followed by a zone-dependent reporting delay. Upon the within-farm mortality exceeding a precomputed fraction of flock capacity (Supplementary Information, Table S4), the farm became suspected of being infected. Enhanced passive surveillance, as occurring within regulated zones (protection and surveillance zones of 3 km and 10 km around detected farms, respectively), reduced the mortality suspicion threshold by 70%, effectively advancing time-to-suspicion by one day for all production types (Supplementary Information, Section 3, Tables S10–S13). Once suspicion was triggered, a reporting delay was sampled from a series of zone-dependent detection delay probabilities, with shorter delays given for farms in regulated zones.

Pre-shipment testing was performed on stage 1 broiler farms located in the high-risk zone (Fig. 1) following initial case detection. Farms were sampled once in the 3–7 days prior to shipment. Detection was simulated through sampling from a binomial probability:

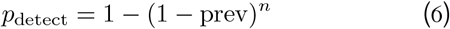

where prev was the within-farm prevalence at the farm’s current day since infection, and *n* was the number of samples drawn (default *n* = 20). Once the within-farm model predicted the onset of mortality (as given by the time-to-first-death having elapsed for that production type), detection probability was set to 1 on the assumption that infected dead birds would be preferentially sampled.

Contact tracing, akin to shipment-mediated transmission, operated through the broiler supply chain and was initiated upon detection of an infected stage 2 broiler farm. Sibling farms (those that received shipments from the same stage 1 source on the same day) were identified and, if infected, scheduled for detection after a short delay. Uninfected sibling farms (as would occur when stage 2 broiler infections were not shipment-mediated) were placed under enhanced surveillance, whether in a regulated zone or not.

All preventively-culled farms underwent routine testing. If a farm was infected but not detected at the time of culling, post-culling detection (following a sampled delay) was assumed to be certain given that birds either found dead or exhibiting signs consistent with HPAI would be prioritized for testing. Detection probabilities and delay distributions are provided in Supplementary Information (Section 3, Tables S10–S13).

#### 2.3.4 Intervention strategies

A national movement standstill was activated upon index case detection, where all planned shipments were paused for a three-day period. Production cycles of active farms were accordingly extended by this duration, under the simplifying assumption that farms would absorb the cost of additional holding time without any flock turnover. Given the short duration, this seemed a reasonable assumption.

Further interventions were implemented through period-dependent policy bundles, consisting of reactive and preventive culling, control zones, movement bans, and confinement. A summary of interventions is given in Table 2, with the complete intervention schedule described in the Supplementary Information (Section 4, Table S14).

**Table 2.** Summary of period-dependent control measures. Interventions escalated across the epidemic periods as response capacity increased and targeting was refined. Day ranges are from first detection (day 0), with Period 1 having began seven days before first detection.

| Measure | Period 1<br>*days –7 to 23* | Period 2<br>*days 24–54* | Period 3<br>*days 55–111* |
| --- | --- | --- | --- |
| Reactive cull delay (days from confirmation) | 2 | 1 | 0 |
| Preventive cull radius (km) | 1 | 3 | 3 |
| Preventive cull targets | All production types | Chickens (broiler 1, broiler 2, layer) | Chickens (broiler 1, broiler 2, layer) |
| Culling capacity | Fixed | Incrementing | Fixed (at maximum) |
| Maximum daily culls (farms/day) | 7 | 60 | 60 |
| National movement standstill | 3 days, upon first detection | — | — |
| Confinement (housing order) | Not active | Broiler 2 chickens, organic ducks (20% susceptibility reduction) | Broiler 2 chickens (20% susceptibility reduction) |

All detected farms underwent culling, with culling ability subject to national capacity constraints. Culling capacity was either fixed (period 1 and 3) or incrementing (period 2) to reflect how human resources were adapted by veterinary services as the epidemics progressed. Preventive culling of undetected farms occurred within a specified radius of detected farms, was targeted to specific farm production types, and was activated upon cumulative incidence exceeding 10 cases. Both reactive and preventive culls competed for the same daily capacity, with reactive culls receiving priority. Culling capacity and strategy (i.e. application radius, daily capacity, and production-type targets) varied by period.

Regulated zones of 3 km (“protection zone”) and 10 km (“surveillance zone”) radii were precomputed for each farm. These zones were used to impose movement bans, enhance surveillance, and restrict restocking upon detection.

Farm types with outdoor production (organic duck and stage 2 broiler farms) were subject to housing orders upon activation of confinement measures. Confinement was assumed to occur immediately, reducing the farm type’s susceptibility *ϕ*_*p*(*j*)_ by 20%. As susceptibility was in both the community and external force of infection terms (Eqs. 1, 3), confinement reduced exposure from both routes.

Both the intensity and scale of interventions changed across the three epidemic periods. In Period 1, limited response capabilities meant a fixed daily culling capacity of 7 farms per day, preventive culling applied indiscriminately to all farm types within 1 km of a detected farm, and prolonged laboratory reporting delays. In Period 2, culling capacity was augmented through variable daily increases to a preset maximum of 60 farms per day, preventive culling zone size was expanded to 3 km with only chicken production types (broiler 1, broiler 2, layer) targeted, and confinement measures were implemented for outdoor farms (broiler 2, organic duck). In Period 3, confinement measures now applied to only broiler 2 farms, and maximum daily culling capacity remained fixed at the maximum reached in Period 2.

Period transitions were triggered at fixed times relative to the start of the simulation (initial infection), with Periods 1, 2, and 3 beginning on simulation days 1, 32, and 63, respectively.

Transitions were spaced roughly four weeks apart so that each in-simulation period matched the four-week duration of the corresponding player phase, and were fixed in advance instead of being contingent on the state of the epidemic at a given time. Results however are reported relative to the first detection, since that was the point from which players observed the epidemic. Because the first detection occurred stochastically, the interval between initial infection and first detection varied across realisations, and thus the period boundaries fell on different days in each realisation when counted from first detection. In the realized epidemic the first detection occurred on simulation day 8, so when observed in terms of days from first detection, Period 1 begins on day *−*7, Period 2 on day 24, and Period 3 on day 55.

#### 2.3.5 Stochastic simulations, model outputs, and trajectory selection

Each stochastic realisation was initialized with a single random number seed that governed every stochastic step in the simulation, starting with selection of the index farm. The index farm was randomly drawn among those in the southern area of the island that met the criteria for production type (broiler 1 or 2), risk score ( *≥*0.005), and remaining production cycle activity at time of infection ( *≥*14 days). All other farms were susceptible if active, or for farms in a downtime period, susceptible upon activity re-initiation. All model events (infection, infectiousness, suspicion, detection, culling) were recorded with their associated characteristics (e.g. event day, infection route, detection or culling method, zone membership, event-associated delay), enabling full reconstruction of the epidemic network and trajectory.

Model calibration proceeded through a series of stages. Within-farm parameters were set to reproduce, for each production type, within-farm trajectories that reflected the short, concentrated infectious period and the rapid rise to a high within-flock prevalence and mortality that are characteristic of HPAI H5-like infection. Relative infectivity and susceptibility parameters were then used to establish the desired influence of each production type in shaping between-farm epidemic dynamics, specifically fixing stage 2 broiler farms as the dominant contributor to overall epidemic spread, as production type has been previously identified as an important determinant of a farm’s contribution to HPAI epidemic dynamics (Andronico et al., 2019). The timing of control measures, implementation delays, and capacity constraints were then configured to produce the desired durations within the three-period framework. Lastly, a structured grid of baseline maximum between-farm transmission rate (*β*_max_) values (that informed only community-based transmission, the primary infection driver) spanning 0.049 to 0.052 was explored. As *β*_max_ principally controlled epidemic size while variations between seeds governed epidemic trajectory shape and timing, *β*_max_ values were chosen to yield epidemics of the targeted size, and shape and timing was left to vary by seed.

The inclusion criteria against which trajectories were filtered were: spatially-heterogenous exponential spread mostly contained in the south of the island prior to propagating northward; an epidemic peak occurring early in period 2; a total epidemic size between 450 and 750 infected farms (roughly 5–8% of the total susceptible population); a long right tail with epidemic extinction prior to 120 days after first detection; and infections represented among all species and production types with transmission driven predominantly by stage 2 broiler farms.

For each *β*_max_ configuration (4 values), multiple stochastic simulations were run via an array of random number seeds. Twenty seeds satisfying the inclusion criteria were retained per *β*_max_ configuration, yielding 80 candidate trajectories.

The Challenge organizing committee (i.e. the authors of this paper) selected a single final trajectory as one that fell within a representative range of the candidate ensemble, which was used for all subsequent data generation, story-boarding, and participant evaluation.

## 3. Results

### 3.1 General epidemic dynamics

Infection dynamics varied among the 80 retained stochastic realisations, with daily detected incidence rising from sporadic detections to peak rates of 16–43 farms per day, occurring 21– 37 days after first detection (Fig. 3). The movement standstill, activated immediately upon first detection, briefly suppressed shipment-based transmission; however, a mean of 17 (range: 9–32) farms had already been infected by the time of its application. The transition to Period 2 occurred 19–24 days after first detection, shortly prior to the peak of the epidemic. By the transition to Period 3 (beginning 50–55 days after first detection) daily incidence had been in decline for all realisations. Sporadic detections (defined as periods where at least 80% of days over a 7-day window recorded no more than one detection) began on average 74 days after first detection, and the last outbreaks were detected on average 87 (70–107) days following first detection.

**Figure 3.**
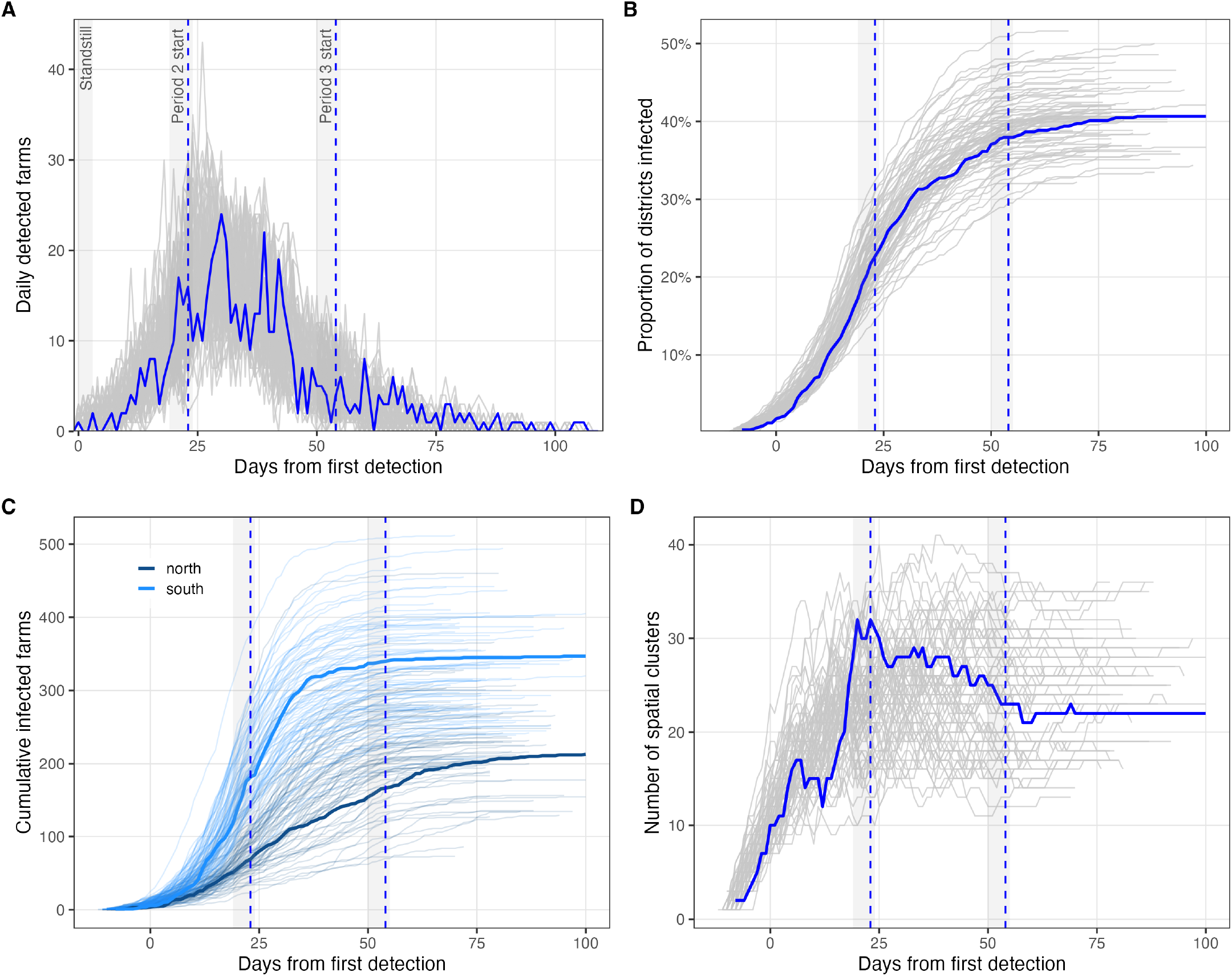
Temporal and spatial dynamics of epidemic spread across conserved stochastic realisations (*n* = 80). In all panels, each thin line (grey in A, B and D, coloured by region in C) is an individual realisation and the thick line (blue in A, B and D, thick coloured lines in C) is the final selected trajectory; the *x*-axis shows days from first detection, which varied between realisations. Grey shaded vertical bands indicate the range of period-transition timing across realisations (arising from variation in the day of first detection), and blue dashed lines indicate the period-transition days of the selected trajectory. Daily detected incidence. The leftmost shaded band marks the 3-day movement standstill after first detection. In all realisations, detected incidence rapidly increased following first detection, peaked on average 28 (range: 21–37) days later, and gradually declined to only sporadic cases in the Period 3 tail. Cumulative spatial coverage expressed as the proportion of districts infected since first detection, indicating early near-exponential spread followed by a gradual plateauing. **(C)** Cumulative infected farms stratified by northern and southern island region membership, indicating a predominance of spread in the southern region during Period 1. **(D)** Number of spatial clusters of infected farms over time, where clusters are groups of infected farms located within 10 km of at least one other infected farm, corresponding to the surveillance zone radius as defined in the model. Clustering indicates the degree of spatial fragmentation, which helps to distinguish local transmission (cluster expansion and merging) from long-distance spread (formation of new clusters); cluster counts therefore increase as transmission events over 10 km occur, and decrease as clusters enlarge and absorb other clusters.

The spatial extent of the epidemic varied among realisations while respecting the trajectory-inclusion constraints previously described. By the end of the simulation, outbreaks covered a mean of 40% (range: 32–52%) of all districts, with the selected trajectory reflecting the average spatial dispersion. In all realisations, the southern DPPA was affected first and accumulated cases more rapidly than the north, consistent with index case location and higher production density of more susceptible and infectious production types (broiler chickens).

Cluster analysis using a 10 km threshold (corresponding to the surveillance zone radius) revealed an early peak in the number of distinct affected areas, reflecting transmission driven by external introductions and long-range animal movements. Outward local spread which grew epidemic size then resulted in a progressive merging and stabilisation of the number of clusters. The per-district infection probability across all realisations was near-certain (>0.95) in the southern DPPA, elevated (>0.75) in the northern DPPA, and low but non-negligible elsewhere (Supplementary information, Section 5, Figure S2).

### 3.2 Selected trajectory

The selected trajectory contained 560 infected farms and ran over the course of 118 days, from first infection to end of final cull. The initial case occurred in the southern DPPA with the majority of infections occurring near the end of Period 1 and beginning of Period 2. Spatial progression with true infection timing is shown by period in Figure 4.

**Figure 4.**
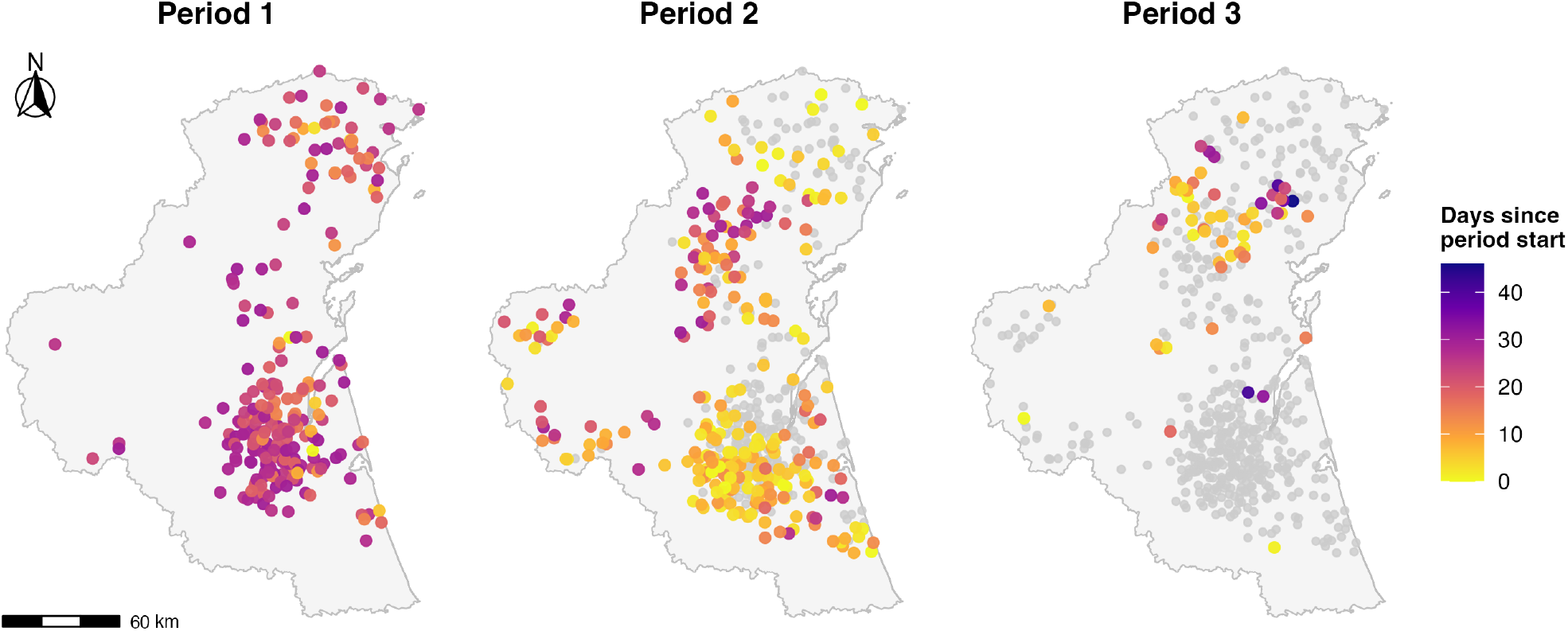
Spatial progression of the selected-trajectory epidemic by epidemic period. Each panel shows the infected farms of the selected trajectory, coloured by day of infection relative to the start of that period (lighter = earlier in the period). Farms infected in previous periods are shown in grey, so the cumulative epidemic is displayed with only current-period infections highlighted. The southern dense poultry production area (DPPA) containing the index case saturates during Period 1 and early Period 2, after which the majority of infections occur in the northern DPPA

Stage 2 broilers accounted for the largest share of outbreaks (*n* = 255, 46%). Following were stage 1 broiler (101, 18%), conventional duck (78, 14%), layer (64, 11%), and organic duck (62, 11%) farms. Community transmission accounted for the vast majority of infections (515, 92%), with external introductions (28, 5%) and network transmission (17, 3%) accounting for the remainder.

A total of 1,188 culls were performed, of which 497 were reactive and 691 were preventive. Passive surveillance was the dominant detection pathway (468, 84%), followed by post-culling detection (74, 13%), pre-shipment testing (14, 2.5%), and contact tracing (4, 0.7%). All infected farms were eventually detected. Culling demand was not uniformly met during the epidemic, with culling demand during Period 1 reaching the national capacity of 7 farms per day for the final 8 consecutive days of the period (Fig. 5). The culling backlog was relieved through Period 2, as capacity was augmented up to a maximum of 60 farms per day. Saturation ceased around the epidemic peak, after which the daily demand remained within the expanded capacity for the remainder of the epidemic.

**Figure 5.**
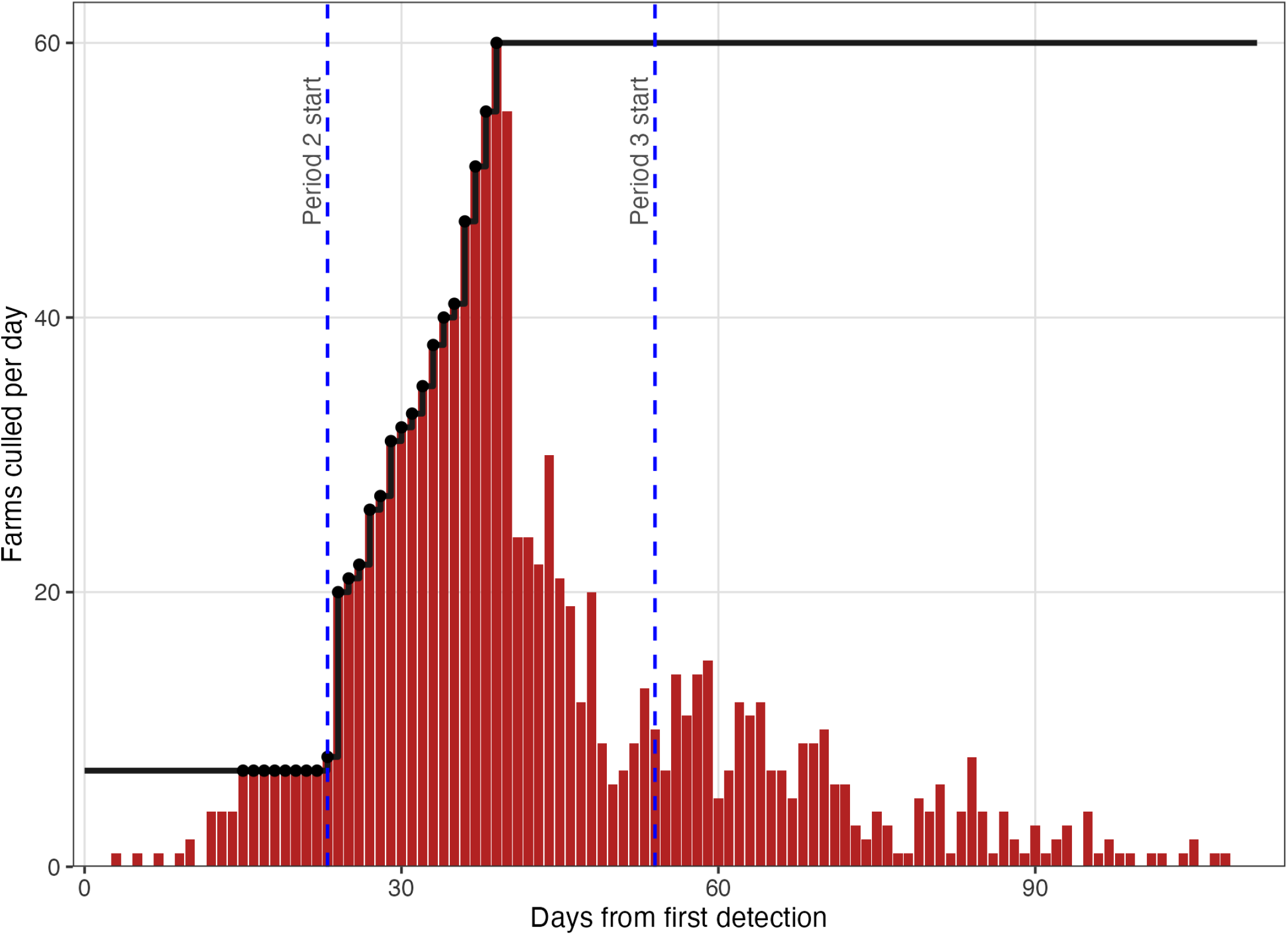
Figure 5. Daily culling activity versus national culling capacity for selected trajectory. Bars indicate farms culled (both reactively and preventively) per day; the black line indicates national culling capacity, which was held at a fixed ceiling of 7 farms per day through Period 1 before being augmented from the start of Period 2 until a maximum capacity of 60 farms per day was reached. Black dots indicate days in which culling capacity was saturated. Culling was saturated for a continuous span of 25 days from day 15–39 after first detection, with 8 of those saturation days occurring in Period 1.

### 3.3 Data provided to participating teams

Prior to Challenge kick-off, teams were provided with the complete synthetic population data, to allow familiarisation and facilitate motivation. This included a farm registry containing the geographic coordinates, county and district of belonging, species, production type, and flock capacity of every farm; a movement dataset detailing broiler supply chain shipments; a farm activity database detailing planned active production periods for all farms; and shapefiles containing Jolly Island administrative boundaries (counties and districts), freshwater features, and the high-risk zone. Mock epidemiological data, in the form of line-lists of detected outbreaks and preventive culls, were also provided, along with an accompanying data-description text describing the structure and contents of each provided file. All population datasets are appended as supplementary material.

All files were hosted on a common server and remained accessible throughout the duration of the Challenge. Access to datasets for subsequent phases was restricted until the start of the corresponding phase.

#### 3.3.1 Surveillance outputs

At the start of each phase, teams were provided four datasets as comma-separated value files: a line-list of outbreaks, a line-list of preventive culls, movement records for the broiler supply chain, and activity periods per farm. The outbreak registry contained farm numeric identifiers, dates of suspicion (if applicable) and confirmation, detection method, culling start and end dates, and cull states (completed, in-progress, or scheduled). The preventive cull list contained the farm identifier, cull start and end dates, and cull states. Though real epidemic data sets often contain overlapping and sometimes conflicting information (e.g. coordinates, administrative location, species presence, or movement dates), for simplicity and cleanliness, conflicting information was intentionally avoided and teams were relied on to cross-reference the demographic datasets to obtain desired farm characteristics. Additionally, in the first phase, teams were provided with three handwritten within-farm mortality ledgers to serve as a window into within-farm dynamics. All phase-explicit datasets are provided as supplementary material.

#### 3.3.2 Situation reports and context

The data release of each phase was accompanied by a narrative situation report. This dated report described the qualitative epidemiological situation, interventions that had been implemented or modified, and a series of questions for which the teams were instructed to provide their responses in the form of a report. The situation reports for each phase can be found in the phase-specific supplementary material.

#### 3.3.3 Participant missions

Each phase opened with descriptive epidemiology questions asking teams to summarise the observed epidemic through a breakdown of outbreaks by species and production type, an incidence timeline, and a spatial representation of detected outbreaks.

Forecasting questions were posed in Phase 1 and Phase 2, asking teams to predict the temporal and spatial evolution of the epidemic (by farm type, if possible) over the subsequent four weeks, assuming the active management strategy had remained unchanged. No forecast was requested in Phase 3 as by then the epidemic had resolved.

Parameter inference was introduced in Phase 1. Through a question asking teams to characterize the relative contribution of galliform versus palmiped farms to viral spread, teams had to move beyond description and towards attributing transmission events to specific elements of the population.

Counterfactual policy evaluation formed the largest and most demanding category, expanding across subsequent phases as the interventions evolved. In Phase 1, teams were asked to determine the effect of restricting preventive culling to chicken farms only during the second epidemic period, and the tradeoff value between maintaining preventive culling efforts or reducing the reactive culling delay by one day. In Phase 2 this latter question was retained, with the addition of also assessing the consequences of relaxing confinement protocols for duck and all farms during the third epidemic period, and of prohibiting restocking for all farms within surveillance zones (as opposed to only those that were culled). In Phase 3, as the epidemic had now subsided, teams were asked when depopulated farms could be safely restocked without risking epidemic flare-up, how many outbreaks the preventive culling strategy had averted, and what daily culling capacity would have been necessary at the end of Period 1 to have enabled epidemic fade-out by the end of Period 2.

## 4. Discussion

The HPAI Modelling Challenge was the second modelling challenge conducted for an infectious livestock disease, following the ASF Challenge in 2020 (Ezanno et al., 2023), and the first to address highly pathogenic avian influenza. It required the construction of a fully synthetic system, replete with a dynamic poultry population and production supply chain, an epidemic realized upon them, and control strategies that adapted as the epidemic progressed.

As with the ASF Challenge, the aim was not to reproduce a specific real-world HPAI epidemic. Rather, the objective was to generate an epidemic that captured essential features of HPAI virus spread in an intensive poultry production system: distance-dependent local transmission, spread through the supply chain, wildlife-driven introductions, and delayed escalating control interventions. Accordingly, the model was not parameterised to any specific HPAI subtype. Akin to the generative model used for the ASF Challenge, the final model that was used to generate the synthetic data in the HPAI Challenge belonged to a set of any number of model formulations that could have produced representative trajectories (Picault et al., 2022). The specific parameterisation reflected a set of plausible assumptions, though given the number of customisable interacting parameters in the model (*n* = 136), likely there exist multiple alternative parameterisations that could have resulted in similar trajectories, and would be an area for exploration in future instantiations of this initiative.

A distinct feature of the HPAI Challenge was the provision of complete data to participating teams. Unlike the RAPIDD Ebola Challenge, which introduced fog-of-war realism through post-hoc noise, underreporting, and data degradation (Ajelli et al., 2018) or the ASF Challenge which included incomplete population and imperfect case detection data (Picault et al., 2022), here a deliberate choice was made to provide teams with clean and complete population, movement, activity, and case data sets. By having a fully-observed population registry, along with clean and harmonized movement and activity records, teams were able to focus on the core analytical challenges of transmission characterisation, parameter inference, and policy support, rather than on data imputation or noise correction. While real-world farm outbreak data is inherently messy, subject to underreporting, incomplete and patchy population knowledge, and limitations related to data quality (e.g. conflicting information between different data sets), forcing teams to contend with these issues on top of the challenge of constructing a model under tight time constraints was considered beyond the intended scope of the Challenge. Now that several groups have working models, future iterations could deliberately introduce data imperfections, under-detections, missing movement records, reporting delays, and uncertain or mixed farm species compositions, making the handling of these complications an explicit objective.

Within-farm dynamics proceeded deterministically in the model. As farms hold thousands of birds, chance variation in the infection process would likely average out, and a deterministic SEIRD formulation avoided the computational cost of simulating each individual bird or re-solving a stochastic within-farm process every time a farm became infected. With only a small fraction of total farms ever becoming infected, computing independent within-farm trajectories across the whole population would have been unnecessarily expensive and likely produce minimal difference in the resulting between-farm dynamics. By solving the within-farm model once per production type, a mechanistic basis for both the infectiousness profiles and detection timings that drove between farm spread and control was able to be maintained instead of having to assign those qualities statistically.

The infectious period was modelled as Erlang-distributed rather than exponential, as an exponential infectious period implies high variance and unrealistically long infectious tails that can skew transmission in metapopulation systems (Vergu et al., 2010; Wilkinson and Sharkey, 2018). In contrast, an Erlang formulation produces a more-concentrated infectious period that better-represents the within-farm amplification and decline pattern observed in the spread of highly-contagious viral diseases (Ferguson et al., 2001). In combination with the time-since-infection function that mapped within-farm prevalence to between-farm infectious pressure, transmission was concentrated within an approximately two-week window (Supplementary Information, Section 5, Figure S3).

An important outcome of the HPAI Modelling Challenge is the creation of a fully observed reproducible simulated HPAI epidemic, providing a benchmark against which modelling teams can evaluate and compare quantitative outbreak response methods beyond the formal event itself. While the player-facing datasets reflect the level of epidemiological details typically seen in high-resource settings, they can be readily adapted to emulate an information-limited context (e.g. making case and population data only available at the district level), a flexibility that is key to supporting the Challenge objective of capacity development across a range of environments. By framing this event as a broad epidemiological challenge, rather than restricting participation to a single task such as forecasting or intervention assessment, teams were expected to address multiple facets of outbreak response: description, prediction, inference, and policy evaluation. This scope choice helped to reduce the barrier to entry for groups with diverse methodological expertise, while fostering capacity development across the full spectrum of activities needed in epidemic response support. Lastly, because the simulated epidemic took place over a roughly 100-day period, the Challenge progressed at a pace approaching that of a real outbreak, and served to expose participants to the time pressures and iterative decision-making that are characteristic of actual outbreak response.

This paper, describing the design, generative model, and data of the Challenge, is the first of a collection. Subsequent contributions to this special issue document the approaches taken by participating teams, and it closes with an analytic paper that provides comparative and ensemble analyses. The accompanying editorial situates the collection as a whole.

## 5. Code availability

Code is available at: https://forge.inrae.fr/gaelbeaunee/hpaimodelling-challenge-2026

## 6. Data availability

Data are available at: https://doi.org/10.5281/zenodo.23069101

## 7. Funding

This work was supported through the WiLiMan-ID project, funded by the European Union via grant agreement 101083833

## Acknowledgement

The authors would like to thank the risk managers and veterinary authorities for their participation in the stakeholders panels informing Challenge design, the WiLiMan-ID project for hosting this initiative, and the team at Biofaction KG for webpage, mail and social media support.

## Notes

https://zenodo.org/records/23069101

### Competing Interest Statement

The authors have declared no competing interest.

