## Supplementary material for "A modelling challenge to improve preparedness for highly pathogenic avian influenza epidemics: objectives, design and synthetic data generation": Supplemntal information

### 1. General information

The model used to generate the synthetic epidemiological data was implemented in R (R Core Team, 2024) with C++ integration via the Rcpp package (Eddelbuettel and Francois, 2011). The simulation engine (`BetweenFarmSim`) is documented in full in the *Technical Process Documentation*, available in the Challenge repository. The model ran in discrete time with daily time steps. Stochastic processes were evaluated at sub-daily resolution ($\Delta t = 0.1$ days) as a $\tau$-leap approximation to continuous-time dynamics. All files required to reproduce the synthetic epidemic are available on the public Challenge repository (see Data availability section of main text).

The epidemic was run across the Jolly Island poultry population, composed of 9,160 farms across five species-specific production types: chicken stage 1 broilers and chicken stage 2 broilers, chicken layers, organic duck farms, and conventional duck farms. In the main text are described the population structure and dynamics (Section 2.2), within-farm compartmental structure (Section 2.3.1), and between-farm transmission equations (Section 2.3.2).

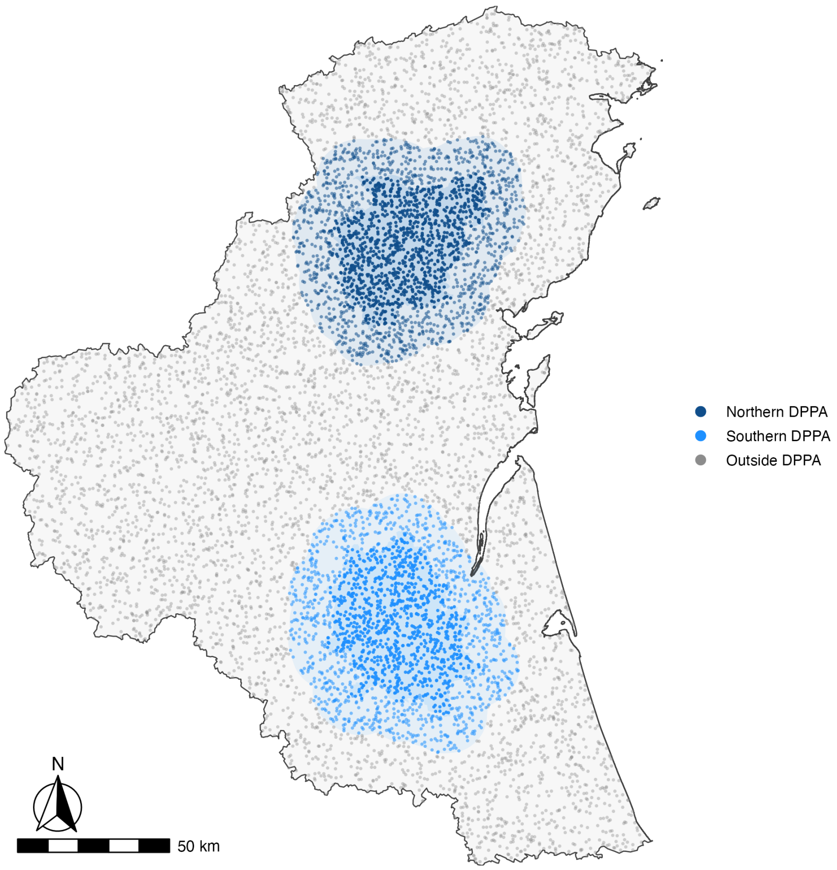


**Figure S1.** Locations of all 9,160 farms coloured by densely populated poultry area (DPPA) membership. Each DPPA consists of a core of contiguous districts plus a transition ring of 12 km. Farm density was 0.66 farms km$^{-2}$ in the northern DPPA core versus 0.52 farms km$^{-2}$ in the southern DPPA core (darker colours), decreasing to 0.39 and 0.33 farms km$^{-2}$ in the respective transition rings. The average density outside these areas was 0.21 farms km$^{-2}$.

### 2. Full model parameterisation

#### 2.1 Structural parameters

Parameters that determine the epidemic simulation structure are provided in Table S1.

**Table S1.** Structural simulation parameters.

| Parameter | Description | Value | Source |
|---|---|---|---|
| `n_days` | Total simulation duration (days) | 120 | Design |
| `dt` | Sub-daily time step for stochastic evaluation (days) | 0.1 | Design |
| `model_start_date` | Simulation calendar start date | 14 Dec 2025 | Design |

#### 2.2 Within-farm parameters

Parameters governing within-farm dynamics are listed in Tables S2–S5. Within-farm transmission rates ($\beta$) were production-type-specific, with smaller broiler and organic duck farms following density-dependent transmission and larger conventional duck and layer farms following frequency-dependent transmission (main manuscript, Section 2.3.1). The infectious period was represented as a chain of 10 Erlang stages to concentrate infectiousness around its mean and reduce prolonged low-level transmission.

**Table S2.** Within-farm transmission rates ($\beta$) by production type. For density-dependent types, $\beta$ has units bird$^{-1}$ d$^{-1}$; for frequency-dependent types, $\beta$ has units d$^{-1}$.

| Production type | Transmission formulation | $\beta$ | Source |
|---|---|---|---|
| Broiler stage 1 | Density-dependent | 0.0018 | Assumed |
| Broiler stage 2 | Density-dependent | 0.0016 | Assumed |
| Conventional duck | Frequency-dependent | 3.9 | Assumed |
| Layer | Frequency-dependent | 3.1 | Assumed |
| Organic duck | Density-dependent | 0.018 | Assumed |

**Table S3.** Species-specific latent period ($1/\varepsilon$), infectious period ($1/\mu_I$), and case fatality rate ($f$). The infectious period is represented as a chain of Erlang stages ($n = 10$, main manuscript Figure 2); the values below represent the mean total infectious period.

| Species | Latent period $1/\varepsilon$ (days) | Mean infectious period $1/\mu_I$ (days) | Case fatality rate $f$ |
|---|---|---|---|
| Chicken | 1.0 | 5.2 | 0.86 |
| Duck | 1.4 | 3.0 | 0.45 |

Source: all values assumed, but kept broadly consistent with infection data for H5 HPAI in domestic poultry.

**Table S4.** Passive surveillance detection thresholds (mortality fraction of flock capacity that triggers suspicion) under standard and enhanced surveillance, by production type.

| Production type | Passive threshold (standard) | Passive threshold (enhanced) | Source |
|---|---|---|---|
| Broiler stage 1 | 0.0052 | 0.00156 | Assumed |
| Broiler stage 2 | 0.0055 | 0.00165 | Assumed |
| Conventional duck | 0.0065 | 0.00195 | Assumed |
| Layer | 0.0030 | 0.00090 | Assumed |
| Organic duck | 0.0150 | 0.00450 | Assumed |

Enhanced threshold = standard threshold $\times$ 0.3. Enhanced surveillance is triggered by membership of a regulated zone.

**Table S5.** Within-farm model outputs by production type, used as inputs for the between-farm simulation. All times expressed as mean days since farm infection. Suspicion is triggered when daily mortality exceeds its production type-specific threshold (Table S4).

| Production type | Time to infectiousness | Time to first death | Time to suspicion (passive) | Time to suspicion (enhanced) |
|---|---|---|---|---|
| Broiler stage 1 | 1 | 5 | 6 | 5 |
| Broiler stage 2 | 1 | 5 | 7 | 6 |
| Conventional duck | 2 | 5 | 9 | 8 |
| Layer | 1 | 6 | 9 | 8 |
| Organic duck | 1 | 4 | 5 | 4 |

### 2.3 Between-farm parameters

Between-farm transmission parameters are given in Tables S6–S9. Transmission kernel ($K_{ij}$) parameters are listed in Table S6 along with the global between-farm transmission rate $\beta_{\max}$. The between-farm force of infection was also modulated by production type infectivity ($\psi$) and susceptibility ($\phi$) modifiers for infectious and susceptible farms respectively (Table S7), and by the within-farm infectious proportion as a function of time since infectiousness.

**Table S6.** Global between-farm transmission rate and spatial transmission kernel parameters.

| Parameter | Description | Value | Source |
|---|---|---|---|
| $\beta_{\max}$ | Maximum baseline community transmission rate ($d^{-1}$) | 0.049 | Calibrated |
| $h_0$ | Height (maximum value) | 1 | Fixed |
| $r_0$ | Scale (distance at which the kernel decreases to half of $h_0$) | 8 | Calibrated |
| $\alpha$ | Shape (governs rate of decay) | 3 | Calibrated |
| $d_{\max}$ | Maximum transmission distance cutoff (km) | 30 | Assumed |

$\beta_{\max}$ was selected from a structured grid spanning 0.049–0.052. The value shown is that of the selected trajectory.

**Table S7.** Production type-specific infectivity ($\psi$) and susceptibility ($\phi$) modifiers for between-farm transmission, used to calibrate the relative contribution of each production type to overall epidemic propagation. Broiler stage 2 farms served as reference ($\phi = \psi = 1.0$).

| Production type | Infectivity modifier $\psi$ | Susceptibility modifier $\phi$ | Source |
|---|---|---|---|
| Broiler stage 1 | 0.85 | 0.91 | Calibrated |
| Broiler stage 2 | 1.00 | 1.00 | Reference |
| Conventional duck | 0.09 | 0.06 | Calibrated |
| Layer | 0.52 | 0.43 | Calibrated |
| Organic duck | 0.28 | 0.09 | Calibrated |

**Table S8.** External (wildlife-driven) daily transmission rate ($\beta_q^{\text{ext}}$) by epidemic period.

| Epidemic period | Simulation days (days from first detection) | $\beta_q^{\text{ext}}$ | Source |
|---|---|---|---|
| Period 1 | 1–31 (−7 to 23) | 0.004 | Calibrated |
| Period 2 | 32–62 (24–54) | 0.001 | Calibrated |
| Period 3 | 63–119 (55–111) | 0.000 | Design |

Days in parentheses are counted from first detection in the selected trajectory, which occurred on simulation day 8 (observed day 0).
External transmission intensity decreased across periods to reflect the reduction in transmission opportunities associated with the end of the migration season. No external transmission occurred in Period 3, ensuring epidemic extinction.

**Table S9.** Additional between-farm transmission parameters.

| Parameter | Description | Value | Source |
|---|---|---|---|
| $m_q^{\text{det}}$ | Value taken by the infectiousness multiplier $m_i(t)$ (main manuscript, Eq. 1) once farm $i$ is suspected or detected, in epidemic period $q$. | 0.5 (Period 1), 0.25 (Period 2), 0.125 (Period 3) | Assumed |
| $r_j$ | Farm-level relative risk of exposure to infected wild birds, given as a logistic function of the distance from freshwater bodies. | Median $2.7 \times 10^{-5}$ (IQR $2.3 \times 10^{-8}$, $1.1 \times 10^{-2}$) | Derived |

### 3. Detection probability and delay distributions

Detection was modelled through four pathways: passive surveillance (mortality-driven suspicion), pre-shipment testing of stage 1 broiler farms in the high-risk zone, contact tracing through the broiler supply chain, and post-culling detection. Passive surveillance relied on a production-type-specific mortality threshold (Table S4). All subsequently described processes are governed by the following delay distributions.

**Table S10.** Reporting delay distribution following passive suspicion, by control zone membership. Values are unnormalised sampling weights; one delay is drawn per farm with probability proportional to its weight.

| Control zone membership | 1 day | 2 days | 3 days | 4 days | 5 days | Note |
|---|---|---|---|---|---|---|
| Out of zone | — | 0.55 | 0.45 | 0.30 | 0.10 | Longer delays; standard passive surveillance |
| In zone | 0.40 | 0.40 | — | — | — | Shorter delays; enhanced surveillance within control zone |

Weights were normalised at time of sampling. '—' indicates the delay is not available for that context.

**Table S11.** Culling start delay distribution. Values are unnormalised sampling weights; one delay is drawn per culling event with probability proportional to its listed weight.

| 0 days | 1 day | 2 days | 3 days | 4 days | 5 days | 6 days | 7 days | Source |
|---|---|---|---|---|---|---|---|---|
| 0.200 | 0.350 | 0.350 | 0.150 | 0.080 | 0.010 | 0.005 | 0.001 | Assumed |

Weights were normalised at time of sampling. Applies to both reactive and preventive culling. Reactive culling delay is measured from confirmation; preventive culling delay from the decision to cull. The delay reflects time required for team and equipment preparation, travel, and setup.

**Table S12.** Post-culling detection delay distribution for infected farms not yet detected at time of culling. Values are unnormalised sampling weights; one delay is drawn per farm with probability proportional to its listed weight.

| 2 days | 3 days | 4 days | 5 days | 6 days | 7 days | Source |
|---|---|---|---|---|---|---|
| 0.20 | 0.30 | 0.25 | 0.15 | 0.07 | 0.03 | Assumed |

Weights were normalised internally at time of sampling. Detection probability was set to 1 at the time of culling for all infected farms, reflecting preferential sampling of already-dead or symptomatic birds. The delay reflects time required for sampling, processing, and reporting.

**Table S13.** Contact tracing detection delay distribution for infected sibling farms identified upon detection of a stage 2 broiler farm. Values are unnormalised sampling weights; one delay is drawn per farm with probability proportional to its listed weight.

| 1 day | 2 days | 3 days | 4 days | Source |
|---|---|---|---|---|
| 0.20 | 0.40 | 0.30 | 0.10 | Assumed |

Weights were normalised internally at time of sampling. Sibling farms sharing a stage 1 broiler source on the same shipment day were identified upon detection of any stage 2 farm. Uninfected siblings were placed under enhanced surveillance rather than scheduled for detection.

### 4. Complete intervention schedule

Interventions were implemented through period-dependent policy bundles. Table S14 gives the complete parameterisation of all control measures by epidemic period. Changes in intervention parameters across periods are described in the main manuscript (Section 2.3.4).

**Table S14.** Complete intervention schedule by epidemic period. Days are counted from first detection; Period 1 opens on day −7 and period transitions occurred at days 24 and 55.

| Parameter / measure | Period 1 (days −7 to 23) | Period 2 (days 24–54) | Period 3 (days 55–111) |
|---|---|---|---|
| **Reactive culling** | | | |
| Additional reactive cull delay (days from confirmation to cull start) | 2 | 1 | 0 |
| **Preventive culling** | | | |
| Cumulative incidence trigger (no. detected farms) | 10 | 0 (immediate) | 0 (immediate) |
| Preventive cull radius (km) | 1 | 3 | 3 |
| Targeted production types | All (broiler 1, broiler 2, conventional duck, layer, organic duck) | Broiler 1, broiler 2, layer | Broiler 1, broiler 2, layer |
| **Culling capacity** | | | |
| Capacity rule | Fixed | Ramp | Fixed |
| Maximum daily culls (farms) | 7 | 60 | 60 |
| Daily ramp increment (farms/day) | — | 4 | — |
| **Movement restrictions** | | | |
| National standstill (days, upon first detection) | 3 | — | — |
| Movement ban radius around detected farms (km) | 10 | 10 | 10 |
| **Confinement** | | | |
| Enabled | No | Yes | Yes |
| Targeted production types | — | Broiler stage 2, organic duck | Broiler stage 2 |
| Susceptibility multiplier | — | 0.8 | 0.8 |
| **Zone restocking ban** | | | |
| Enabled | No | No | No |
| **Control zones (all periods)** | | | |
| Protection zone radius (km) | 3 | 3 | 3 |
| Surveillance zone radius (km) | 10 | 10 | 10 |

Reactive and preventive culls competed for the same daily culling capacity. Period 2 culling capacity increased incrementally from the preceding day's capacity, where the daily increment was drawn from a Poisson distribution with mean of 4 farms per day, to a ceiling of 60. Confinement reduced susceptibility by the multiplier shown. Zone radii were precomputed for each farm and remained constant across all periods. Zone membership affected surveillance thresholds, reporting delays, and movement bans.

### 5. Additional model dynamics

Two additional descriptive outputs of the model are provided here: the spatial distribution of infection risk across the candidate trajectories (Figure S2), and the production-type-specific within-farm trajectories that underlie the between-farm simulation (Figure S3).

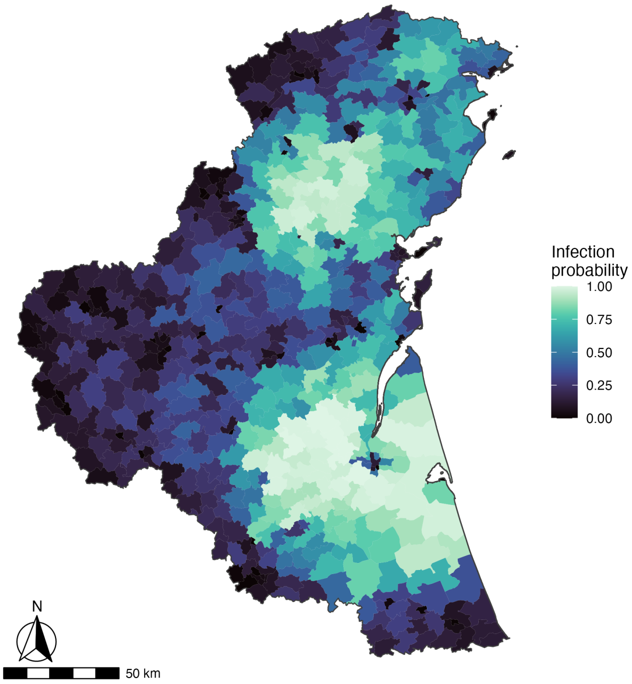


**Figure S2.** Per-district infection probability across the 80 candidate trajectories, defined as the proportion of realisations in which at least one farm in the district was infected. Districts in the two high-density zones were near-certain to have outbreaks, with districts outside these areas (i.e. western region) less likely but still susceptible, reflecting long-distance movement-network transmission.

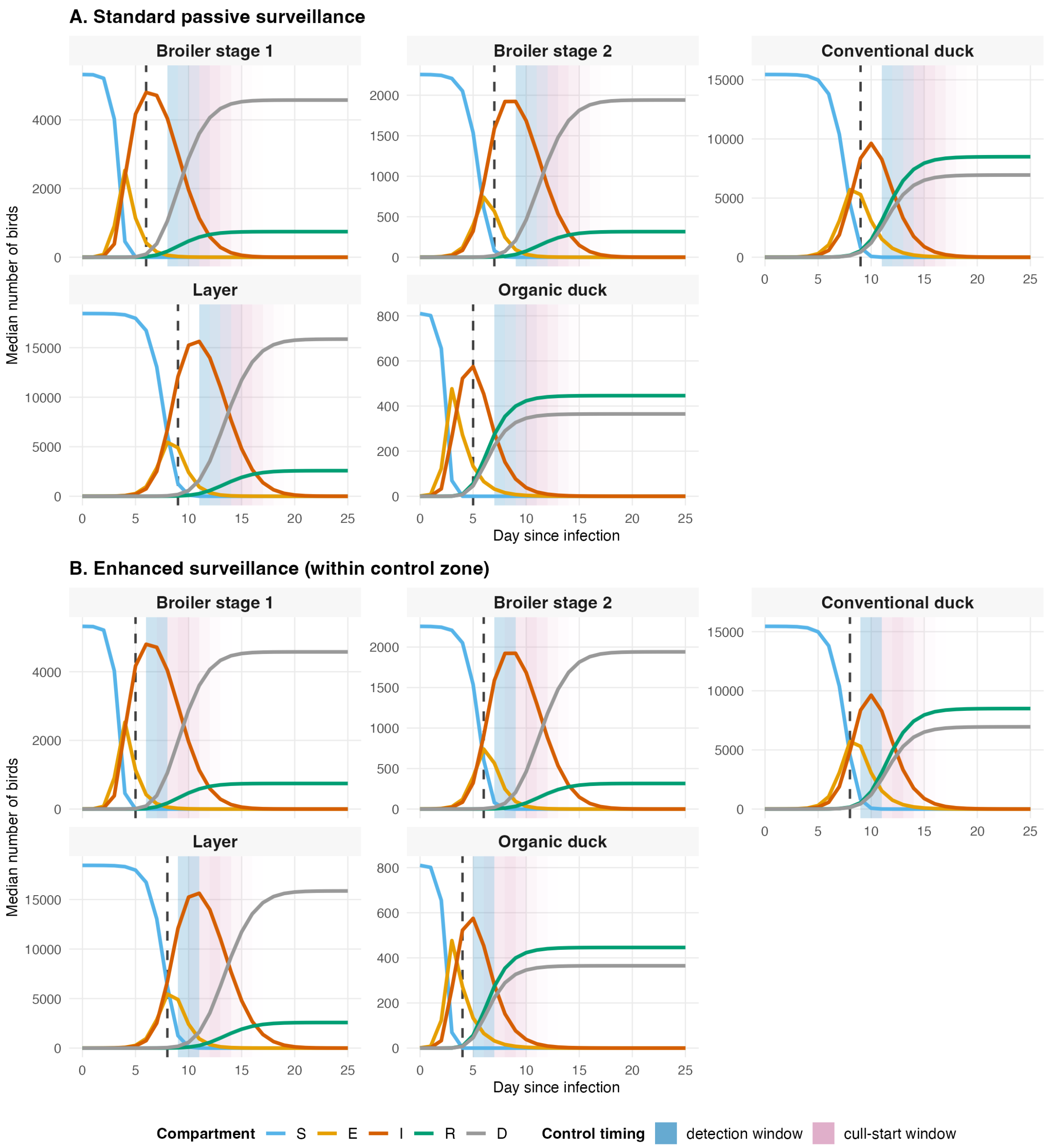


**Figure S3.** Median within-farm SEIRD trajectories by production type with overlain detection and culling timing windows under standard **(A)** and enhanced **(B)** passive surveillance. Each panel shows the trajectory for the median farm size of that production type, expressed as absolute quantity of birds in each compartment (S, susceptible; E, exposed; I, infectious; R, recovered; D, dead) as a function of day since infection. Shaded bands indicate the window over which detection and culling would be expected to occur, and were derived by convolving the timing of the suspicion trigger with the associated delay distribution (Tables S10–S13). In panel B, the narrowing and left-shift of the windows relative to panel A reflects the lower detection threshold under enhanced surveillance. Differences in $y$-axis scale between chicken broiler stage 1, stage 2, and organic duck farms compared to chicken layer and conventional duck farms arise because the former cohort followed density-dependent transmission, while frequency-dependent transmission was used for the latter.

### 6. Situation reports and phase data

Three situation reports and phase-specific datasets were provided to participating teams at the start of each Challenge phase. Each report described the current epidemiological situation, summarised the active control measures, and provided a set of questions for response. The reports were dated to align with the phase data release dates.

The three situation reports are available on the public Challenge repository (see Data availability section of main text).
